# ZHX2 alleviates vascular remodeling and smooth muscle cell proliferation by transcriptional regulation of GADD45G

**DOI:** 10.1101/2023.10.15.562431

**Authors:** Siyuan Fan, Liuye Yang, Xuelian Wu, Yichen Wu, Pengchao Wang, Baoru Qiao, Yue Li, Kai Huang, Zhe Zheng

## Abstract

**Objective:** This study explores the role of ZHX2 in vascular remodeling, specifically focusing on its effects on VSMC proliferation, migration, and neointima formation following vascular injury. Methods and Results: Data from both human atherosclerotic samples and a mouse carotid injury model indicated a decrease in ZHX2 levels. *In vivo*, ZHX2 overexpression reduced neointima formation in mice subjected to carotid artery ligation. *In vitro*, ZHX2 inhibited the proliferation and migration of primary VSMCs. Conversely, specific knockout of ZHX2 in SMCs *in vivo* or knockdown of ZHX2 in primary VSMCs had opposite effects. RNA-seq analysis revealed that ZHX2 overexpression significantly affected the expression of cell cycle-related genes. Using Chromatin Immunoprecipitation Sequencing (ChIP-seq) and luciferase reporter assays, we demonstrated that ZHX2 plays a crucial role in the transcriptional regulation of GADD45G, identifying GADD45G as the downstream target responsible for ZHX2-mediated effects. Conclusions: ZHX2 emerges as a key player in pathological vascular remodeling, suppressing VSMC proliferation and migration through its regulatory impact on GADD45G transcription and cell cycle-related gene expression.

## Introduction

Vascular Smooth Muscle Cells (VSMCs), primarily found in the media layer of blood vessels, are crucial for controlling the dilation and constriction of these vessels (1). VSMCs are inactive in the arteries of healthy adults. These smooth muscle cells play important roles in maintaining vascular tone and integrity, regulating luminal pressure, and distributing blood volume. However, VSMCs are not static, and they retain remarkable plasticity. When blood vessels are exposed to certain stimuli, such as inflammation and aging, VSMCs become activated, leading to enhanced proliferation, migration, and synthesis ability, ultimately promoting blood vessel remodeling (2). They play a crucial role in vascular remodeling and maintaining vascular homeostasis by supporting the structural adjustment of existing blood vessels and the formation of new ones (3). Vascular remodeling is a complex process that involves both beneficial and maladaptive responses to various stimuli, which might lead to chronic cardiovascular disease (4). Vascular remodeling can either be beneficial or pathological (5). A common feature of pathological vascular remodeling is neointimal hyperplasia (6, 7), which is an important process in several vascular proliferative diseases, including atherosclerosis (8, 9), hemodialysis vascular access dysfunction (10), hypertension (11), pulmonary arterial hypertension (12), restenosis after angioplasty (13, 14), and venous bypass graft stenosis (15). Under these pathological conditions, activated VSMCs proliferate, migrate excessively, and accumulate under the endothelium (16). The new layer they form between the lumen and the internal elastic lamina is called neointima. An Understanding of how VSMCs change from differentiation to proliferation may contribute to the prevention and treatment of not only restenosis after percutaneous coronary intervention (PCI) but also a series of pathological vascular diseases characterized by neointimal hyperplasia remodeling-related diseases.

The zinc fingers and homeobox (ZHX) family includes ZHX1, ZHX2, and ZHX3, and these proteins have similar unique structures, containing two C2H2-type zinc finger motifs and four or five HOX-like homeodomains (17). Previous studies have suggested that ZHXs can function as positive (18) or negative transcriptional regulators (19). ZHX2 plays the role of oncogene or tumour suppressor gene in different tumours. For example, in hepatocellular sarcoma, ZHX2 can inhibit the expression of the oncogenic gene alpha-fetoprotein (AFP) (20). A number of results showed that in malignant tumours such as thyroid cancer (21), chronic lymphocytic leukemia (22, 23), and multiple myeloma (24, 25), the expression of ZHX2 was negatively correlated with the prognosis of appealing patients. Suggesting that ZHX2 may play the role of a tumour suppressor gene in patients with the above tumours. However, ZHX2 promotes tumorigenesis in clear cell renal cell carcinoma (26) and breast cancer (27). Genome-wide association studies (GWAS) studies show that ZHX2 is associated with intima-media thickness (IMT) (28-30). However, the role of ZHX2 in VSMC remains unclear.

This study found that the expression of ZHX2 decreased in the carotid artery of mice with CAL and in the neointima of restenotic vessels after PCI in humans. Our research has shown that the overexpression of ZHX2 has the ability to restrict the proliferation of VSMC, which, in turn, inhibits the intimal hyperplasia of CAL model mice. By analyzing the results of RNA-seq and ChIP-seq, we found that ZHX2 binds to the Growth Arrest and DNA-Damage-Inducible 45 Gamma (GADD45G) promoter, inhibiting GADD45G transcription and ultimately impeding VSMC proliferation. Our research has identified a new genetic pathway in the reprogramming of VSMCs, which includes ZHX2 and its target gene, GADD45G. This regulatory network could be a promising target for treating pathological vascular remodeling.

## MATERIALS AND METHODS

### Animal procedures

Animal studies were conducted in compliance with the 8th edition (2011) of the National Institutes of Health Guidelines for the Care and Use of Laboratory Animals. Animal care and experimental procedures received approval from the Institutional Animal Care and Use Committee (IACUC) at Huazhong University of Science and Technology (IACUC number 3445). The ZHX2 Flox/Flox (ZHX2 fl/fl) mice were obtained from Shanghai Biomodel Organism Science & Technology Development Co., Ltd., and the tamoxifen-inducible transgenic mice (Myh11-CreERT2) was from The Jackson Laboratory (Stock No. 019079). Female ZHX2 fl/fl mice were mated with male Myh11-CreERT2 mice to produce female ZHX2 fl/+ and male ZHX2 fl/+/Myh11-CreERT2 offspring. These offspring were then crossbred to yield male ZHX2 fl/fl/Myh11-CreERT2 or ZHX2 +/+/Myh11-CreERT2 mice. Male ZHX2 fl/fl/Myh11-CreERT2 mice were crossed with their female ZHX2 fl/fl littermates to generate the experimental ZHX2 fl/fl/Myh11-CreERT2 mice, and ZHX2 +/+/Myh11-CreERT2 mice were mated with their female ZHX2 +/+ littermates to produce the control ZHX2 +/+/Myh11- CreERT2 mice. Using block randomization, animals were randomly allocated to the experimental and control groups. To induce ZHX2 deficiency, mice at the age of 6 weeks were injected with 50 mg/kg of tamoxifen consecutively for five days. The control mice also underwent the same tamoxifen treatment. Genotyping was carried out using Polymerase Chain Reaction (PCR). Fig S2.

### Immunohistochemistry and immunofluorescence staining

The freshly harvested tissues are fixed immediately in 4% paraformaldehyde (PFA) for 24 hours at 4°C. After fixation, the samples are embedded in paraffin and sectioned into 6- micrometre-thick slices. These slices are then stained with Hematoxylin and Eosin (H&E) and immunofluorescence (IF). To prepare for staining, the sections are first deparaffinized and rehydrated. Antigen retrieval is performed by boiling the samples in citrate buffer (pH 6.0) at 95°C for 20 minutes. The slices are then permeabilized by incubating in PBS containing 1% Triton X-100 for 15 minutes. To block non-specific binding, the sections are sealed with goat serum for one hour at room temperature. The primary antibodies are diluted in the blocking buffer according to the manufacturer’s instructions. The slides, coated with the primary antibodies, are incubated overnight at 4°C in a humidified chamber. Subsequently, the samples are stained with fluorescence-labeled secondary antibodies, and the cell nuclei are marked with DAPI. Imaging is performed using an Olympus fluorescence microscope, and ImageJ software is utilized for a blind measurement of either the intensity of the IF signal or the percentage of cells positive for the indicated staining. The primary antibodies were used as follows: ZHX2 (Proteintech, 20136-1-ap), αSMA (Proteintech, 67735-1-Ig), KI67 (Proteintech, 27309-1-AP).

### RT-qPCR

We extracted total RNA from rat cells and tissues employing the TRIzol reagent (D9108A, TaKaRa Bio). The RNA was then reverse-transcribed into complementary DNA (cDNA) using the RR036A PrimeScript™ RT Reagent Kit (Perfect Real Time) supplied by TaKaRa. The amplification products were quantified using SYBR Green (Vazyme), utilizing a PRISM 7900 Sequence Detector System (Applied Biosystems, Foster City). For the quantification of all genes, we used either GAPDH or β-actin as the internal control. The primer sequences used are provided in Supplementary Table 1.

### Cell Culture and Treatment

Primary smooth muscle cells were prepared from SD rats, weighing between 150-180g, using an enzymatic digestion method. The aorta of the rats was surgically removed under aseptic conditions, and the external membrane was then stripped under a dissecting microscope. The aorta was cut into fragments of about 1-2mm and digested for 2 hours in an enzyme solution containing collagenase type II (3mg/mL; C6885, 700.3U/mg, Sigma). and elastase (1mg/mL, E1250, 6U/mg, Sigma). The cells were subsequently cultured in SMCM medium and cultured in a humidified atmosphere containing 5% CO2 at 37°C with experiments conducted from the 3rd to 6th generation.

The mice primary VSMCs were isolated from the aorta of 10-week-old C57BL/6J male mice via an enzymatic digestion method. The mouse aorta was excised under sterile conditions, washed in PBS to remove fat and connective tissue, and then incubated for 10 minutes at 37°C in 1mL DMEM containing type II collagenase (3mg/mL; C6885, 700.3U/mg, Sigma). The outer membrane of the aorta was subsequently stripped, and the tissue was minced and transferred into cell culture flasks. It was then incubated for 30 min in solutions of type II collagenase and for 60 min in elastase (1mg/mL, E1250, 6U/mg, Sigma), respectively. Digestion was halted, and the harvested cells were placed in SMCM medium and cultured in a humidified atmosphere containing 5% CO2 at 37°C. HEK 293T cells, supplied by Procell Biotech (Wuhan, China), were cultured in a 5% CO2 incubator at 37°C. The culture medium used was DMEM supplemented with 10% FBS.

### Western Blot

Cells or tissues were lysed using pre-chilled lysis buffer supplemented with protease inhibitors. Subsequently, the protein concentration was determined by employing the BCA method. Equal quantities of protein were combined with sample buffer, followed by SDS-PAGE electrophoresis and immunoblotting. The specific antibodies utilized are listed below: ZHX1 (Proteintech, 13903-1-AP), ZHX2 (Proteintech, 20136-1-AP), ZHX3 (Proteintech, 29397-1-AP), CCND1 (Proteintech, 26939-1-AP), PCNA (Proteintech, 10205-2-AP), αSMA (Proteintech, 14395-1-AP), E2F1 (CST, #3742), GADD45G (ABclonal, A10286), GAPDH (Proteintech, 60004-1-Ig), αTUBULIN (Proteintech, 80762-1-RR).

### Carotid Artery Wire Ligation Injury Model

All surgeries were performed under sterile conditions. The establishment of the model used a blind method. Both the experiment operators and statistical analyzers did not participate in the prior tamoxifen treatment. Male C57BL/6 mice, aged eight weeks, were anesthetized using an intraperitoneal injection of pentobarbital sodium. For the ligation group, the left common carotid artery was carefully dissected from the surrounding tissue under a microscope and ligated using a 6-0 silk ligature. For the control group, a sham surgery was performed involving the dissection, but not ligation, of the right common carotid artery. For localized adenovirus delivery, Ad-Control or Ad-ZHX2 (4×10 pfu) were blindly packaged by 70 μL Pluronic gel F-127 (Keygen, China) to extend virus contact time and delivery to the carotid arteries immediately after ligation, respectively. Following a period of 14 days, mice were humanely euthanized, and the common carotid arteries were surgically removed. These arteries were subsequently fixed with 4% formaldehyde, embedded in paraffin, and cross-sectioned. The sections were then stained using hematoxylin and eosin (H&E) and Masson’s trichrome to observe elastin and Immunofluorescence.

### Recombinant adenovirus production

Adenoviral vectors, engineered to either express control or carry the ZHX2 coding DNA sequence (cDNA), were used for *in vitro* experiments involving vascular smooth muscle cells (VSMCs). In parallel, similar adenoviral vectors designed to express either scramble short hairpin RNA (shRNA) or ZHX2-targeted shRNA was also employed. For *in vivo* studies, these modified adenoviral constructs were administered to infect the carotid arteries directly.

### EdU

After undergoing the specified treatment for a duration of 12 hours, vascular smooth muscle cells (VSMCs) were subsequently plated into 96-well plates at a density of 3 × 10^4 cells per well. Upon completion of the plating process, the cells were subjected to culture. Following the culture period, a medium containing EdU was introduced to the cells for an additional 2 hours. Subsequent to this incubation, the cells were immobilized using a 4% paraformaldehyde solution. The EdU Incorporation Assay was then conducted in accordance with the guidelines provided by the manufacturer (Beyotime, C0075S). To visualize and document the results, images were acquired employing the Olympus fluorescence microscope.

### Transwell

Following a 24-hour period of the specified treatment, vascular smooth muscle cells (VSMCs) were subjected to digestion and subsequently suspended in a serum-free medium. A cell suspension of 5×10^4 cells (100 μL) was introduced into the upper compartments of transwell culture plates. In parallel, 500 μL of medium containing 10% FBS was added to the lower compartments of the same plates. The entire assembly was then incubated at a temperature of 37°C with a 5% CO2 atmosphere for a duration of 24 hours. After the incubation period, the cells located on the upper surface of the polycarbonate films were delicately eliminated using moistened cotton swabs. Subsequently, the polycarbonate films were cautiously detached from the upper chambers. To proceed, the cells were fixed with pre-chilled methanol for 30 minutes. Following fixation, a staining step was conducted using a 0.1% solution of crystal violet for 15 minutes. The stained cells were rinsed thrice with PBS and subjected to microscopic observation.

### Wound healing assay

Vascular smooth muscle cells (VSMCs) were initially cultured in a 6-well plate, followed by infection and treatment as per the experimental requirements. After a duration of 24 hours, a linear scratch was created across the surface of the cell monolayer using a pipette tip. Subsequently, a medium containing 2% FBS was introduced to the culture. Microscopic imaging of the scratched cell area was performed immediately (0 hours) and following a 24- hour interval. This allowed for visualizing the migration dynamics of the cells. The evaluation of cell migration capacity was carried out by analyzing the degree of healing observed in the scratched region between the initial and final imaging time points.

### RNA-seq

RNA extraction, library preparation, sequencing, and analysis. Total RNAs were extracted using TRIzol Reagent (Invitrogen, cat. NO15596026) following the methods by Chomczynski et al. (DOI:10.1006/abio.1987.9999). DNA digestion was carried out after RNA extraction by DNaseI. RNA quality was determined by examining A260/A280 with NanodropTM OneCspectrophotometer (Thermo Fisher Scientific Inc). RNA Integrity was confirmed by 1.5% agarose gel electrophoresis. Qualified RNAs were finally quantified by Qubit3.0 with QubitTMRNA Broad Range Assay kit (Life Technologies, Q10210). 2 μg total RNAs were used for stranded RNA sequencing library preparation using KCTMStranded mRNA Library Prep Kit for Illumina^®^ (Catalog NO. DR08402, Wuhan Seqhealth Co., Ltd. China) following the manufacturer’s instruction. PCR products corresponding to 200-500 bps were enriched, quantified, and finally sequenced on a Novaseq 6000 sequencer (Illumina) with a PE150 model. Raw sequencing data was first filtered by Trimmomatic (version 0.36), low-quality reads were discarded, and the reads contaminated with adaptor sequences were trimmed. Clean data were mapped to the reference genome of rat from https://ftp.ensembl.org/pub/ using STAR software (version 2.5.3a) with default parameters. Reads mapped to the exon regions of each gene were counted by featureCounts (Subread-1.5.1; Bioconductor), and then RPKMs were calculated. Genes differentially expressed between groups were identified using the edgeR package (version 3.12.1). A FDR cutoff of 0.05 and a fold-change cutoff of 1 were used to judge the statistical significance of gene expression differences.

### ChIP**-seq**

ChIP assay was performed on VSMC by SeqHealth (Wuhan, China). The tissue/cell was fixed in 1% formaldehyde for 10 min at room temperature, after which 0.125 M glycine was added, and the mixture was sat for 5 min to terminate the crosslinking reaction. The tissue was then collected and frozen in liquid nitrogen. The cells were treated with cell lysis buffer, and the nucleus was collected by centrifuging at 2000g for 5. Then, the nucleus was treated with nucleus lysis buffer and sonicated to fragment chromatin DNA. The 10% lysis sonicated chromatin was stored and named "input", and 90% was used in immunoprecipitation reactions with anti-ZHX2 antibody (Proteintech, 20136-1-AP) and named "IP". The DNA of input and IP was extracted by the phenol-chloroform method. The high-throughput DNA sequencing libraries were prepared by using VAHTS Universal DNA Library Prep Kit for Illumina V3(Catalog NO. ND607, Vazyme). The library products corresponding to 200-500 bps were enriched, quantified and finally sequenced on a Novaseq 6000 sequencer (Illumina) with a PE150 model. Raw sequencing data was first filtered by Trimmomatic (version 0.36), low-quality reads were discarded and the reads contaminated with adaptor sequences were trimmed. The clean reads were used for protein binding site analysis. They were mapped to the reference genome of rat from https://ftp.ensembl.org/pub/ using STAR software (version 2.5.3a) with default parameters. The RSeQC (version 2.6) was used for read distribution analysis. The MACS2 software (Version 2.1.1) was used for peak calling.

### Dual-luciferase assay

The company (TsingkeBiotechnologyCo., Ltd) was commissioned to create synthetic sequences of the GADD45G gene promoter for rat (-2000 to +200) and to incorporate them into the pGL3.0 basic vector. Subsequently, a mutated vector was assembled using the reverse PCR technique. In 293T cells, the pRL-TK plasmid was co-transfected alongside either the empty vector or the ZHX2 overexpression vector and the wild-type or mutant reporter gene vectors. After a span of 12 hours, the Dual-Luciferase Reporter Gene Assay Kit II (Beyotime, RG029) was utilized for measurements as per the instructions provided by the manufacturer.

### ChIP-qPCR

The ChIP-qPCR experiments were carried out according to the manufacturer’s instructions using the ChIP Assay Kit (Beyotime, P2078). In brief, VSMCs were treated with PDGF-bb for 12 hours, with 27 μL of 37% formaldehyde added to each milliliter of the culture medium. This was followed by incubation in formaldehyde at 37°C for 10 minutes to crosslink target proteins and corresponding genomic DNA. After replacing the culture medium with glycine solution from the kit, we prepared pre-cooled PBS containing PMSF and used it to suspend and centrifuge the cells. The cells were then scraped, washed twice, and resuspended in SDS Lysis Buffer with 1 mM PMSF. The mixture was incubated on ice for 10 minutes and then sonicated. Subsequently, 8 μL of 5M NaCl was added to 0.2ml of sonicated sample, mixed thoroughly, and heated at 65°C for 4 hours to de-crosslink the proteins and genomic DNA. DNA extraction was performed next using phenol-chloroform. We then prepared a ChIP Dilution Buffer with 1 mM PMSF and diluted the sonicated sample. Some of the sample was taken out as an input and mixed with Protein A+G Agarose/Salmon Sperm DNA. The sample was centrifuged, and the supernatant was transferred to a new centrifuge tube and incubated with an appropriate amount of primary antibody overnight at 4°C with gentle rotation or shaking. Protein A+G Agarose/Salmon Sperm DNA was added next to precipitate protein or complexes recognized by the primary antibody. After centrifugation, the liquid was discarded, and the precipitate was washed sequentially with wash buffer and finally used for qPCR. The primers used were as follows: forward primer: CGAGCGCAAGTAAAGATTCCC, reverse primer: AAAGGCGAGGTGAAATCTGC.

### Statistics

All data are expressed as the mean ± standard error of the mean and were analyzed using SPSS software. Initially, the Shapiro–Wilk test was utilized to assess the normality of the data. For the comparison of two groups, the Student’s Unpaired t-test was applied when the data were normally distributed with equal variances; for normally distributed data with unequal variances, the t-test with Welch’s Correction was employed; and for data that did not follow a normal distribution, the Mann–Whitney U Test was selected. For comparisons involving three or more groups, One-way Analysis of Variance (ANOVA) was utilized for data that were normally distributed with equal variances, followed by Bonferroni’s Post Hoc Tests for significant differences. For data that were normally distributed but had unequal variances, Welch ANOVA and Dunnett’s T3 Post-Hoc Tests were conducted.

## Result

### ZHX2 is downregulated in proliferating VSMCs both *in vivo* and *in vitro*

To understand ZHX2’s role in vascular remodeling, we analyzed atherosclerotic vascular data GSE43292 from the GEO database. The analysis revealed a reduction in the ZHX family expression in human atherosclerosis specimens compared to the control group (Figure 1A). The mouse carotid artery injury model data GSE70410 corroborated these findings (Figure 1B). Further, we assessed the ZHX family protein level in a mouse Carotid Artery Ligation (CAL) model. After 14 days of ligation, we confirmed the successful establishment of the CAL model through Hematoxylin and Eosin (HE) staining (Figure S1A). Western blot assay results revealed a significant reduction in ZHX2 protein in the CAL mice, while ZHX1 and ZHX3 protein levels remained stable compared to the Sham group (Figure 1C). This suggests a potential key role for ZHX2 in vascular remodeling.

**Fig. 1.**
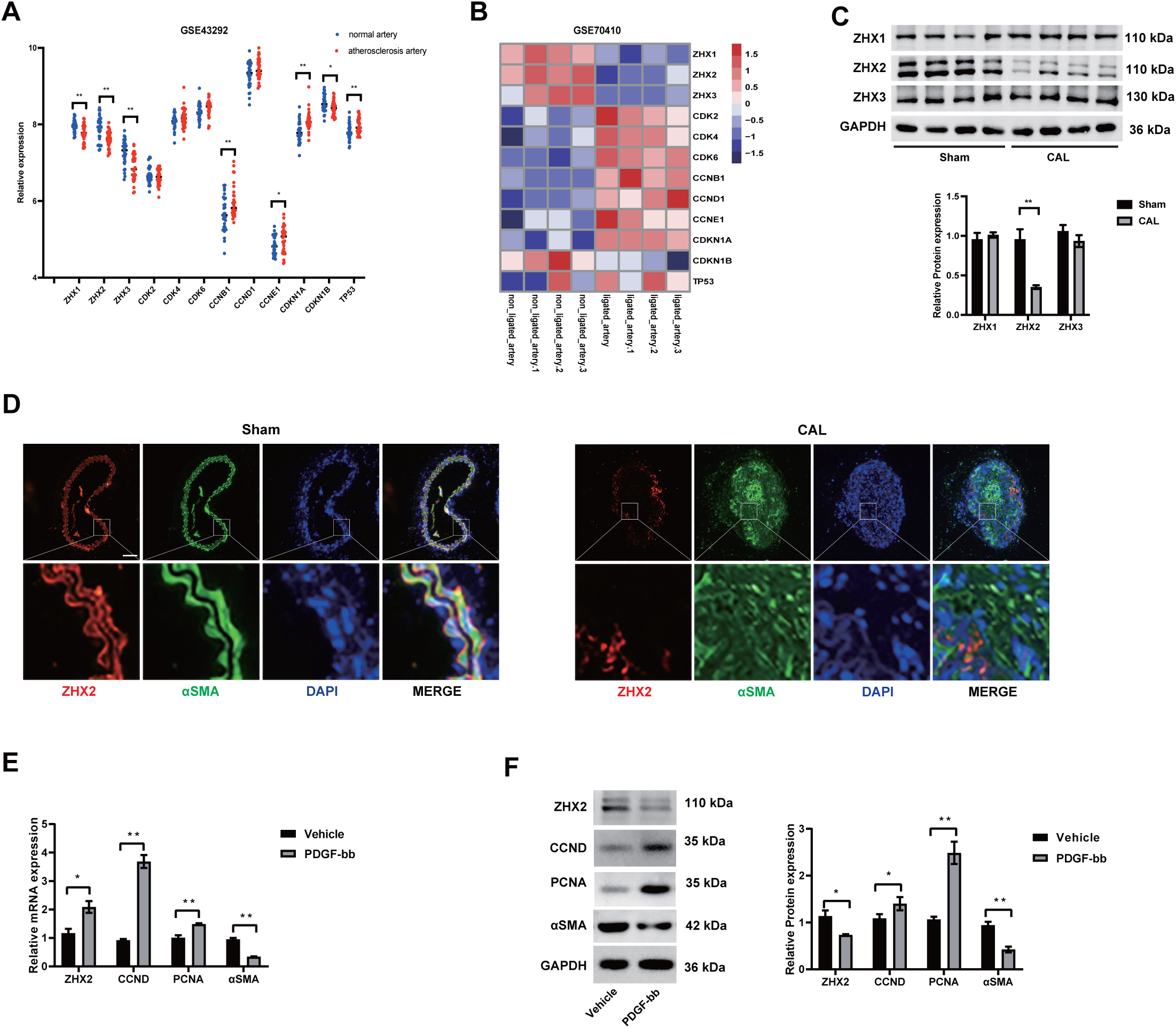
**ZHX2 demonstrates decreased expression levels in both in vitro cultured smooth muscle cells (SMCs) and *in vivo* in atherosclerotic and neointimal SMCs** (A) The GSE43292 atherosclerosis dataset reveals expression levels of both ZHX family genes and genes associated with cell cycle regulation. (B) In the GSE70410 carotid artery injury model dataset, expression of ZHX family genes and cell cycle-related genes is presented. (C) Western blot analysis illustrates the expression of ZHX family proteins in mouse carotid artery injury model samples compared with Sham group samples, accompanied by a statistical graph. (n=4) (D) Immunofluorescence imaging reveals a downregulation of ZHX2 in vascular smooth muscle cells (VSMCs) following a ligation-induced carotid artery injury. Scale bar = 50 μm. (E, F) Both Western blot analysis and real-time quantitative polymerase chain reaction (RT-qPCR) are employed to determine mRNA and protein levels of ZHX2, Cyclin D1 (CCND1), Proliferating Cell Nuclear Antigen (PCNA), and Alpha-Smooth Muscle Actin (αSMA) in VSMCs stimulated by Platelet-Derived Growth Factor BB (PDGF-BB). (n=3) *\*p* < 0.05, *\*\*p* < 0.01, statistical significance determined by Student’s t-test or t-test with Welch’s Correction.

Immunofluorescence results showed a reduction in ZHX2 expression, primarily in αSMA-positive cells, in the CAL mice compared to the Sham group (Figure 1D). To stimulate primary VSMCs *in vitro*, we used PDGF-bb, which caused a decrease in ZHX2 expression at the mRNA level (Figure 1E). The effectiveness of the stimulation was confirmed by PCNA and ACTA2 used as positive controls. Similar results were observed in the Western blot assay (Figure 1F). Collectively, these results hint at a potentially significant role for ZHX2 in vascular remodeling.

### ZHX2 deficiency aggravates neointimal formation induced by ligation and promotes proliferation and migration of VSMCs induced by PDGF-bb

To examine ZHX2’s role *in vivo*, we established a VSMC-specific ZHX2 knockout mouse (Figure S2A). To verify ZHX2’s knockout, we extracted primary mouse vascular smooth muscle cells following regular injections of tamoxifen. Western blot assay results confirmed the knockout of ZHX2 protein in the smooth muscle of CKO mice, compared to Myh11- CreERT2 mice (Figure S2C).

Post the Carotid Artery Ligation (CAL) experiment performed on ZHX2 CKO and Myh11- CreERT2 mice, the tissue sections revealed a thicker intima in the ZHX2 CKO group and the Myh11-CreERT2 group, compared to the Sham group, 14 days after ligation (Figure 2B). To assess smooth muscle cell proliferation during intimal hyperplasia in ZHX2 CKO mice, we evaluated the expression of Ki67 in carotid artery sections with an immunofluorescence experiment. The increased Ki67 expression indicated that ZHX2 knockout could enhance SMC proliferation (Figure 2C).

**Fig. 2.**
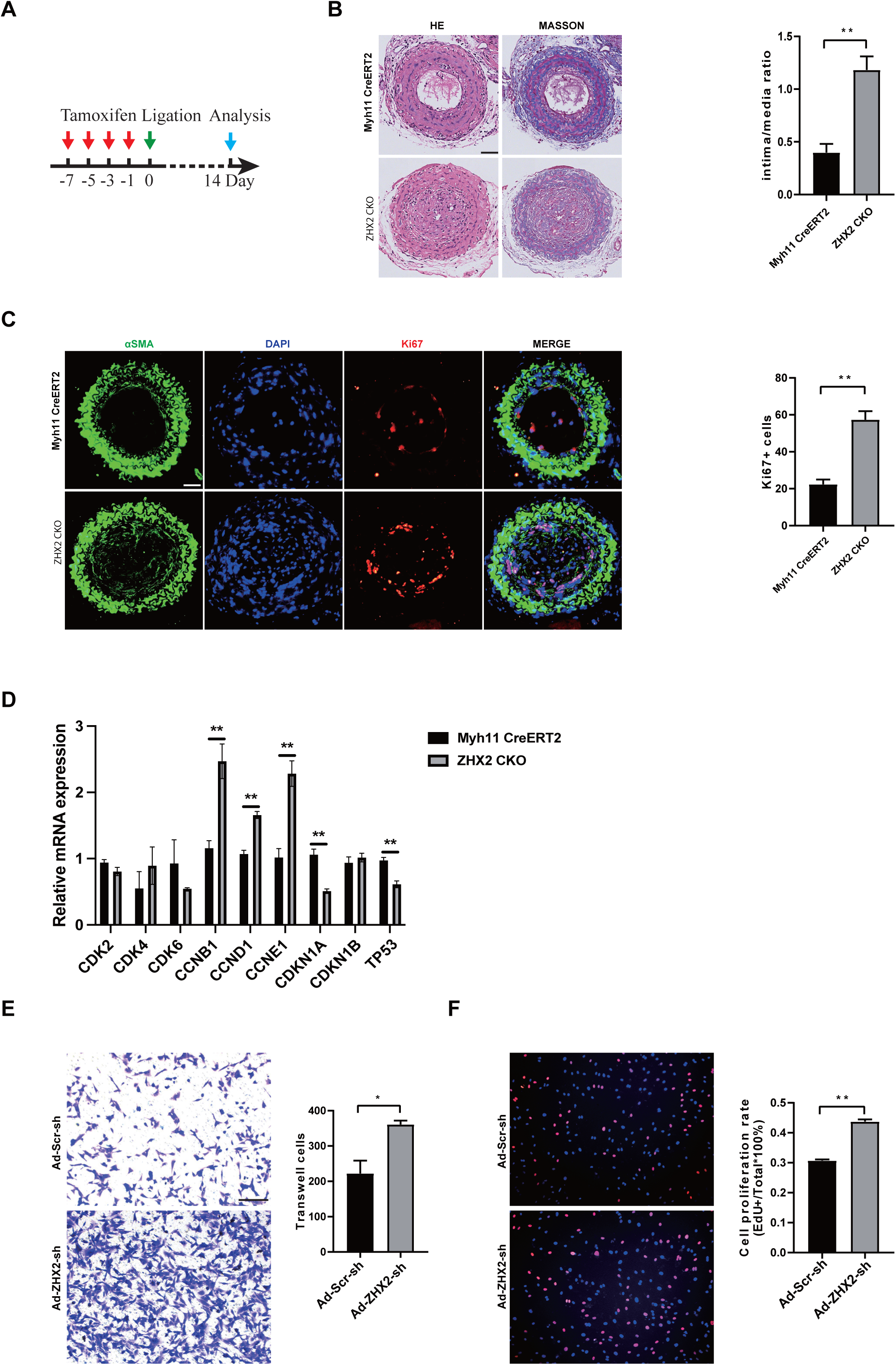
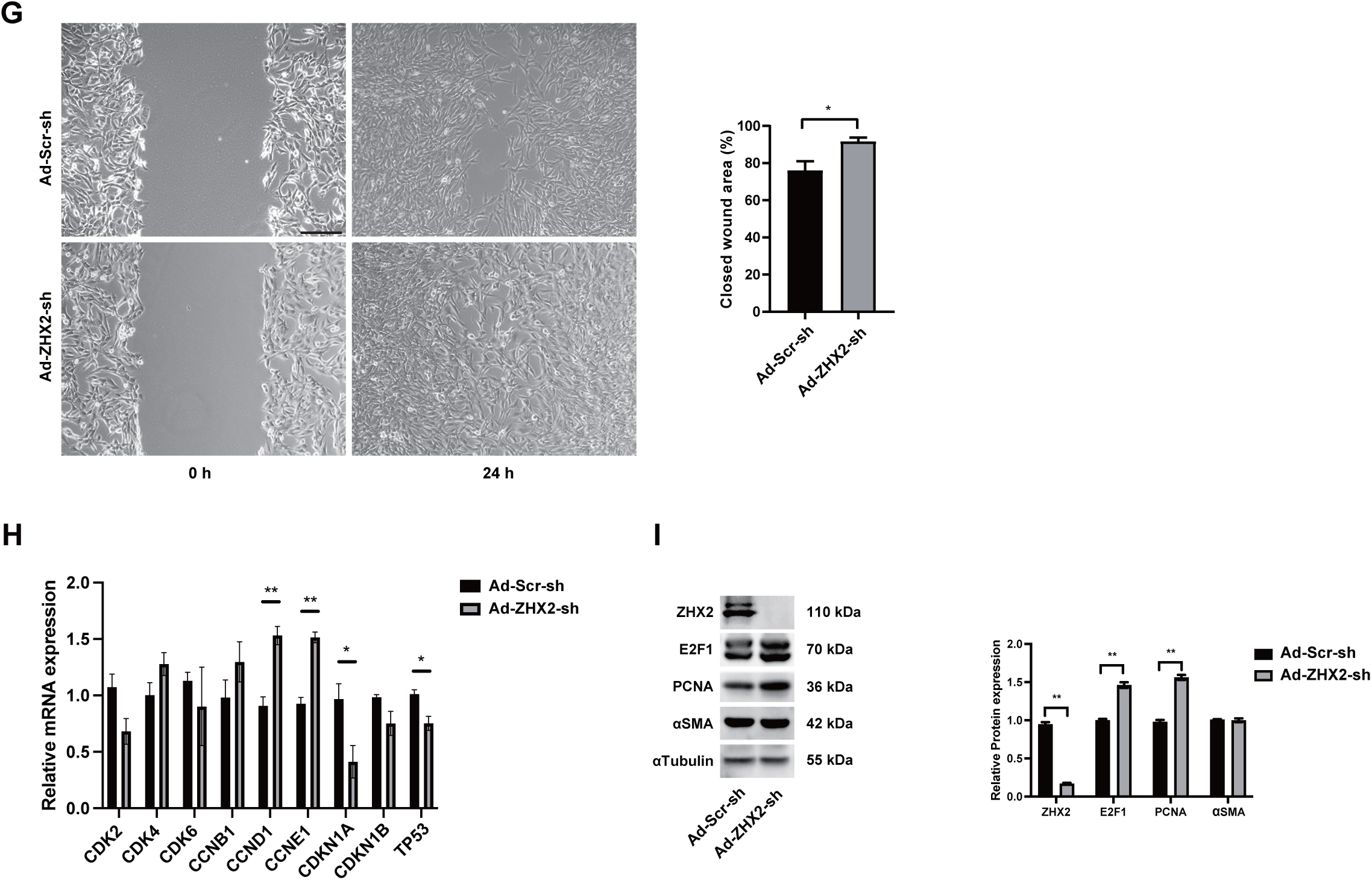
**ZHX2 knockout promotes neointimal formation induced by ligation** (A) Schematic representation of the tamoxifen injection regimen used to induce ZHX2 knockout in VSMCs prior to ligation. (B) Neointimal formation induced by ligation is promoted in VSMCs following specific ZHX2 knockout. (n=3). Scale bar = 50 μm. (C) Immunofluorescence analysis reveals an enhancement in ligation-induced KI67 expression following ZHX2 knockout specific to VSMCs. (n=3). Scale bar = 50 μm. (D) Real-Time Quantitative Polymerase Chain Reaction (RT-qPCR) provides a quantitative analysis of the impact of VSMC-specific ZHX2 knockout on the expression of cell cycle-related genes in a ligation-induced carotid artery injury model. (n=3) (E) A transwell assay demonstrates the effect of ZHX2 knockdown on VSMC migration in *in vitro* cultured smooth muscle cells. (n=3). Scale bar = 100 μm. (F) A 5-Ethynyl-2’-deoxyuridine (EdU) proliferation assay shows the influence of ZHX2 knockdown on VSMC proliferation in *in vitro* cultured smooth muscle cells. (n=3). Scale bar = 100 μm. (G) A wound healing assay reveals the impact of ZHX2 knockdown on VSMC migration in *in vitro* cultured smooth muscle cells. (n=3). Scale bar = 100 μm. (H) RT-qPCR analysis of the effect of ZHX2 knockdown on the mRNA expression levels of cell cycle-related genes in *in vitro* cultured smooth muscle cells. (n=3) (I) Western blot analysis provides quantitative measurement of the impact of ZHX2 knockdown on the protein expression levels of genes associated with cell proliferation and contraction in *in vitro* cultured smooth muscle cells. (n=3) *\*p* < 0.05, *\*\*p* < 0.01, statistical significance determined by Student’s t-test or Welch’s t-test.

Moreover, at the cellular level, primary rat smooth muscle cells infected with either Ad-Scr-sh or Ad-ZHX2-sh were subject to scratch tests, transwell, and EdU experiments. We found that ZHX2 knockdown could promote SMC proliferation and migration (Figures 2E, 2F, 2G). RT-qPCR and Western blot results also suggested that ZHX2 knockdown could enhance SMC proliferation to some extent (Figures 2H, 2I).

In summary, our results suggest that ZHX2 knockdown could promote SMC proliferation both *in vivo* and *in vitro*.

### ZHX2 overexpression alleviates neointimal formation induced by ligation and suppresses proliferation and migration of VSMCs induced by PDGF-bb

Conversely, to determine if ZHX2 overexpression could curb VSMCs proliferation, we established a CAL model in C57BL/6J mice and locally applied adenovirus to overexpress ZHX2. Carotid arteries were collected from the Ad-Control and Ad-ZHX2 groups following ligation. Histological staining of carotid artery tissue sections, 14 days post-ligation, revealed a significant reduction in carotid artery intima thickening in the ZHX2 overexpression group, compared to the control group (Figure 3A). Immunofluorescence of the tissue section showed a decrease in Ki67 positive cells in the Ad-ZHX2 group mice compared to the control group, suggesting that ZHX2 overexpression could inhibit SMC proliferation (Figure 3B). The RT-qPCR assay supported this by indicating that ZHX2 overexpression could restrain SMC proliferation in the CAL model (Figure 3C).

**Fig. 3.**
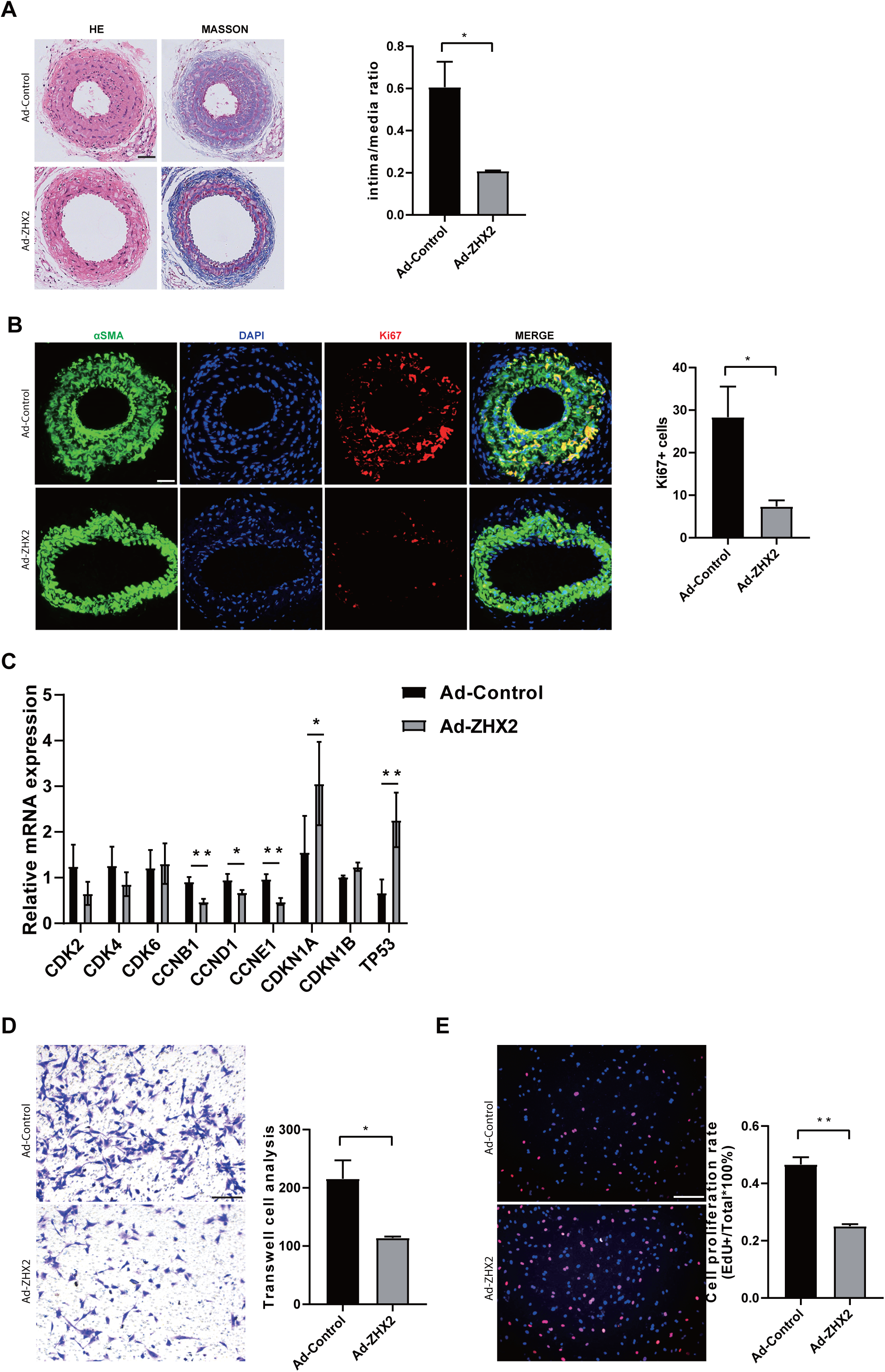
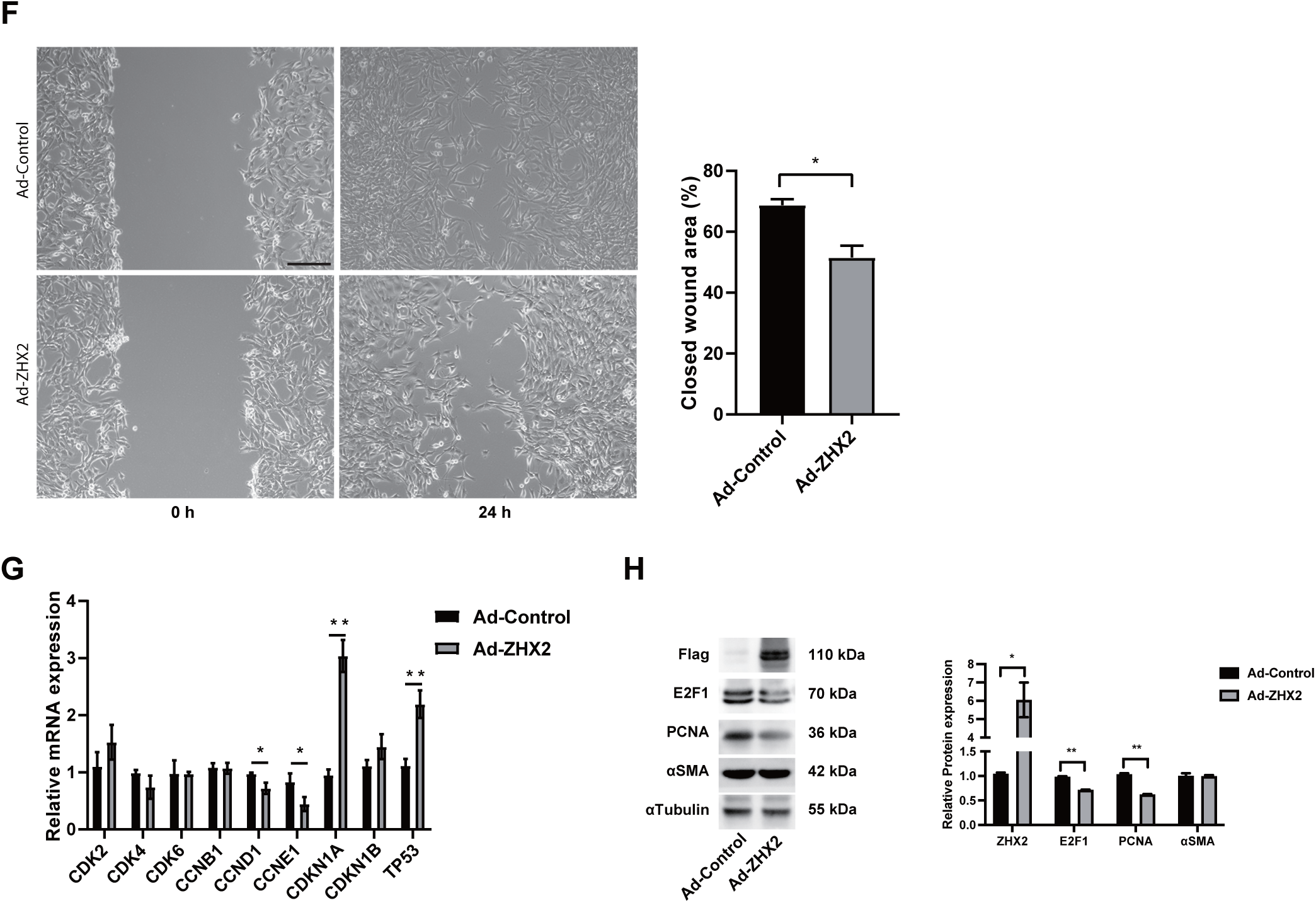
**Overexpression of ZHX2 suppresses ligation-induced neointimal formation.** (A) ZHX2 overexpression inhibits the formation of neointima induced by ligation. (n=3). Scale bar = 50 μm. (B) Immunofluorescence analysis indicates that overexpression of ZHX2 suppresses the expression of the proliferation marker, Ki67, induced by ligation. (n=3). Scale bar = 50 μm. (C) Evaluation of the impact of ZHX2 overexpression on the mRNA expression of cell cycle-associated genes in ligation-induced carotid artery injury, as analyzed via Real-Time Quantitative Polymerase Chain Reaction (RT-qPCR). (n=3) (D) Assessment of the effect of ZHX2 overexpression on VSMC migration in *in vitro* cultured smooth muscle cells, as determined by transwell migration assay. (n=3). Scale bar = 100 μm. (E) Determination of the influence of ZHX2 overexpression on VSMC proliferation in *in vitro* cultured smooth muscle cells, as measured by 5-Ethynyl-2’-deoxyuridine (EdU) proliferation assay. (n=3). Scale bar = 100 μm. (F) Evaluation of the effect of ZHX2 overexpression on VSMC migration in *in vitro* cultured smooth muscle cells, as determined by scratch wound healing assay. (n=3). Scale bar = 100 μm. (G) Analysis of the impact of ZHX2 overexpression on the mRNA expression levels of cell cycle-associated genes in *in vitro* cultured smooth muscle cells, as determined by RT-qPCR. (n=3) (H) Determination of the influence of ZHX2 overexpression on the protein expression levels of genes associated with cell proliferation and contraction in *in vitro* cultured smooth muscle cells, as measured by Western blotting. (n=3) *\*p* < 0.05, *\*\*p* < 0.01, statistical significance determined by Student’s t-test or Welch’s t-test.

At the cellular level, primary rat smooth muscle cells infected with Ad-Control or Ad-ZHX2 demonstrated, through Western blot, successful overexpression of ZHX2 with the use of adenovirus (Figure 3H). Scratch tests, transwell, and EdU experiments results showed that ZHX2 overexpression could inhibit SMC proliferation and migration (Figures 3D, 3E, 3F). RT-qPCR and Western blot assays also revealed ZHX2 overexpression inhibiting SMC proliferation to a certain extent (Figures 3G, 3H).

In summary, our results suggest that ZHX2 overexpression could inhibit SMC proliferation both *in vivo* and *in vitro*.

### RNAseq Discloses Downstream Consequences of ZHX2 in VSMCs Stimulated by PDGF-bb

To further explore the mechanism of ZHX2 inhibiting VSMC proliferation, we used adenovirus to infect primary smooth muscle cells, overexpressed ZHX2 and then conducted RNA-seq to detect the influence of ZHX2 on the transcriptome level of VSMCs. Using RNA-seq heatmap analysis, we evaluated the difference clustering of genes shown by the principal component analysis (PCA) between Ad-Control and Ad-ZHX2 (Figure 4A) and the difference in gene expression shown by the heatmap (Figure 4B). In addition, Gene Set Enrichment Analysis (GSEA) further displayed the co-expression network of cell cycle-related genes after ZHX2 overexpression from the sequencing data (Figure 4C). The GSEA enrichment plot of the most enriched Hallmark gene set "CELL_CYCLE" is shown (Figure 4D). The 10 genes with the highest enrichment scores in the "CELL_CYCLE" gene set are shown in the figure (Figure 4E).

**Fig. 4.**
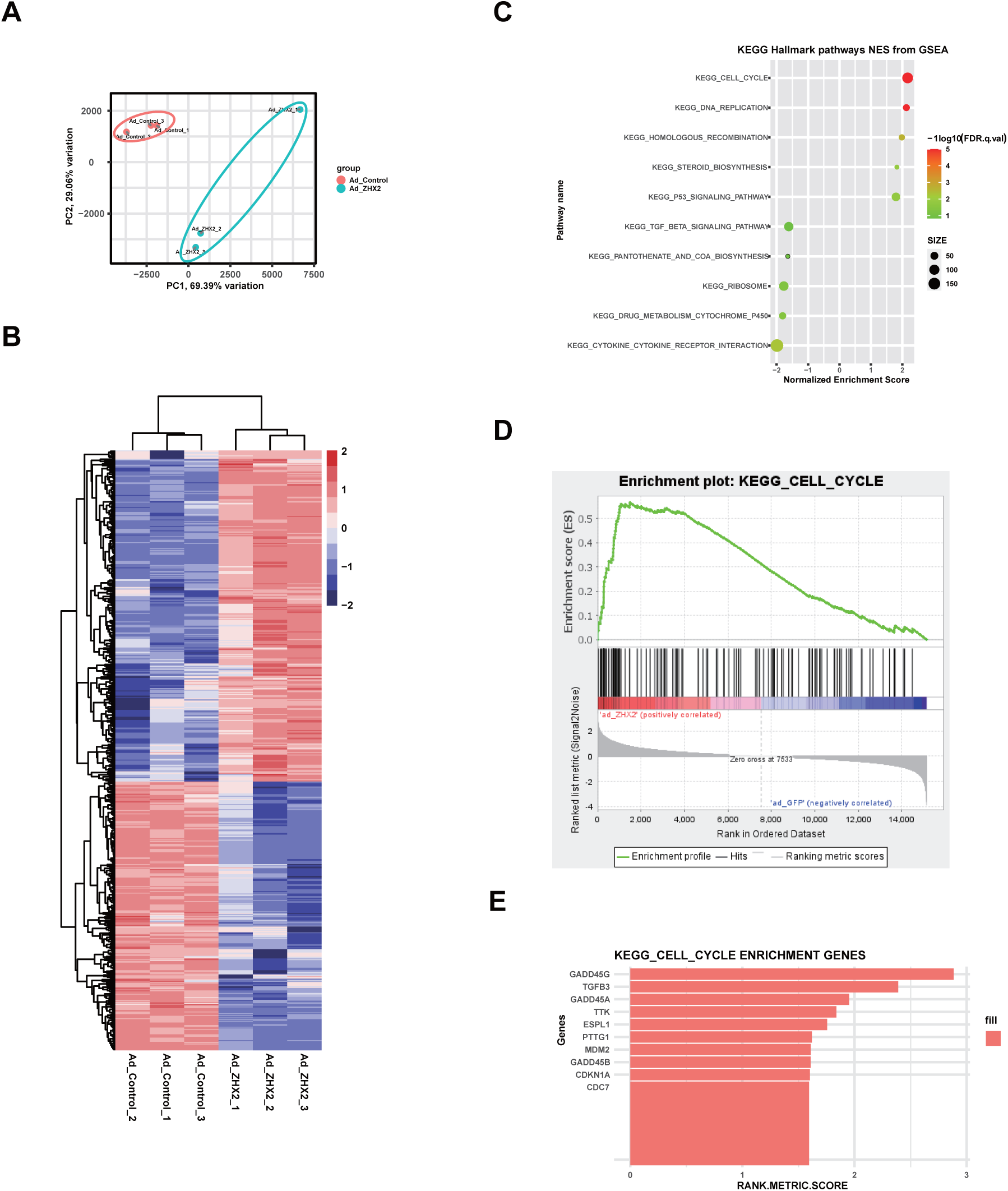
RNA sequencing reveals downstream effects of ZHX2 within Vascular Smooth Muscle Cells (VSMCs) (A) Principal Component Analysis (PCA) illustrates the differential gene clustering in VSMCs treated with Ad-Control or Ad-ZHX2. (B) A heatmap displays the differential gene expression (DEGs) identified from the RNAseq data. (C) Gene Set Enrichment Analysis (GSEA) of genes from VSMCs treated with Ad-ZHX2 versus control, utilizing Kyoto Encyclopedia of Genes and Genomes (KEGG) gene sets for enrichment analysis. (D) A GSEA enrichment plot for the most enriched Hallmark gene set, "CELL_CYCLE", in VSMCs indicates significant enrichment at the transcript level. (E) The top 10 genes with the highest enrichment scores in the "CELL_CYCLE" gene set.

### GADD45G is identified as a direct target of ZHX2

Considering that ZHX2, as a transcription factor, plays a role in transcriptional regulation in various diseases, ChIP-seq were used to explore the target genes regulated by ZHX2. We observed a distinct binding pattern at the transcription start sites (TSS) of differentially expressed genes (Figure 5A). These results suggest that ZHX2 may play a crucial role in the transcriptional regulation of upregulated and downregulated genes. To identify the most likely direct target of ZHX2 that affected the proliferation phenotype of VSMCs, we performed an intersection analysis of differentially expressed genes (DEGs) in RNA-seq, ChIP-seq target genes, and genes in the KEGG Cell Cycle pathway. The analysis showed that GADD45G appears in the intersection of these three datasets (Figure 5B). The interaction of ZHX2 with the promoter of GADD45G through ChIP-seq signal trace images was shown (Figure 5C). Besides, we found that the binding site of ZHX2 with the GADD45G promoter region was highly conserved in humans, rats, and mice (Figure S3B). We constructed GADD45G promoter reporter plasmid and mutated the possible binding sites of ZHX2 (Figure 5D). The luciferase reporter assay showed that ZHX2 overexpression could enhance the transcriptional activity of GADD45G, but had no effect on the promoter with a mutated ZHX2 binding sequence (Figure 5E). The ChIP experiment further confirmed that ZHX2 could directly bind to the GADD45G promoter region (Figure 5F). At the same time, we examined the impact of ZHX2 intervention on GADD45G protein levels and found that ZHX2 knockdown could inhibit the expression of GADD45G protein (Figure 5G), while ZHX2 overexpression could enhance the expression of GADD45G protein (Figure 5H). These results suggested that ZHX2 could directly bind to the promoter of GADD45G.

**Fig. 5.**
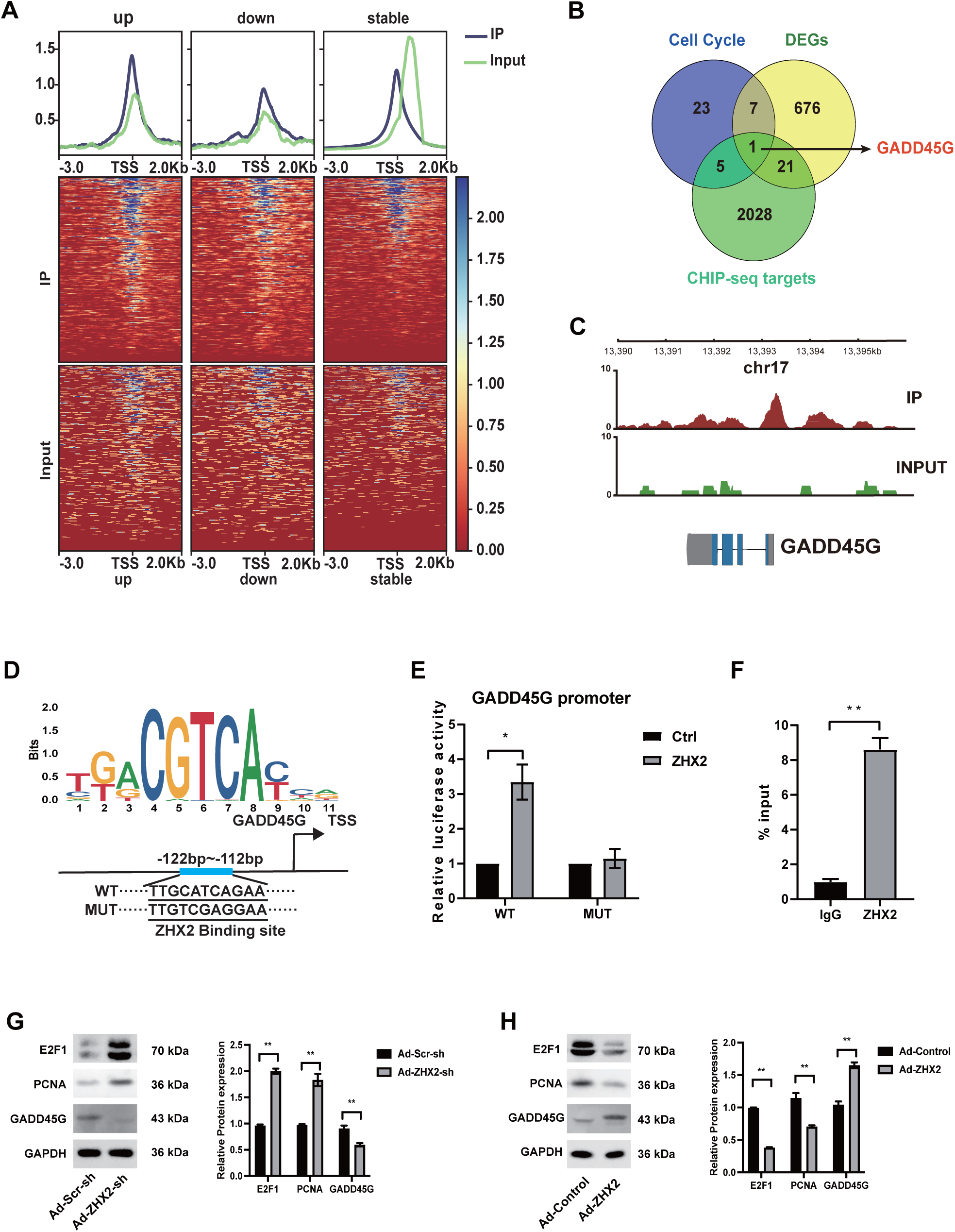
**GADD45G is identified as a direct target of ZHX2** (A) Transcript Start Site (TSS) heatmap showing the binding patterns of upregulated, downregulated, and unchanged genes in Chromatin Immunoprecipitation Sequencing (ChIP-seq) based on RNA sequencing (RNA-seq) data. (B) Venn diagram depicting the overlap of differentially expressed genes (DEGs) from RNA-seq, target genes from ChIP-seq, and genes from the Kyoto Encyclopedia of Genes and Genomes (KEGG) Cell Cycle pathway. (C) ChIP-seq signal track image provides a detailed visualization of the binding intensity within the promoter region of GADD45G, highlighting potential regulatory sites and interactions. (D) Common sequences of ZHX2 identified by Multiple EM for Motif Elicitation (MEME) software (top), putative ZHX2 responsive elements in the GADD45G promoter, and sequences of responsive elements used for constructing the mutant (MUT) vector in the subsequent dual-luciferase reporter gene experiment. (E) Luciferase activity of wild-type (WT) and mutant (Mut) GADD45G promoters in 293T cells overexpressing ZHX2. (n=5) (F) Evaluation of ZHX2 binding to the promoter region of GADD45G in 293T cells by ChIP-qPCR. (n=6) (G, H) Western blot analysis of the effect of ZHX2 knockdown and overexpression on the protein expression levels of GADD45G and proliferation-associated genes in *in vitro* cultured smooth muscle cells. (n=3) *\*p* < 0.05, *\*\*p* < 0.01, statistical significance determined by Student’s t-test or Welch’s t-test.

### Knockdown of GADD45G counteracts the inhibitory effect of ZHX2 on VSMCs

To investigate whether GADD45G mediated the effect of ZHX2 on VSMC proliferation, we conducted transwell and EdU experiments. The results showed that the GADD45G knockdown reversed the effect caused by ZHX2 overexpression (Figures 6A,6B). Subsequent RT-qPCR and Western blot results also validated GADD45G mediated the effect of ZHX2 on VSMC proliferation (Figures 6C, 6D). We also found that GADD45G knockdown could reverse the inhibition of neointima formation caused by ZHX2 overexpression in mice CAL model (Figure 6E). These results indicated that GADD45G knockdown could reverse the inhibition of neointima formation, VSMC proliferation, and migration caused by ZHX2 overexpression.

**Fig. 6.**
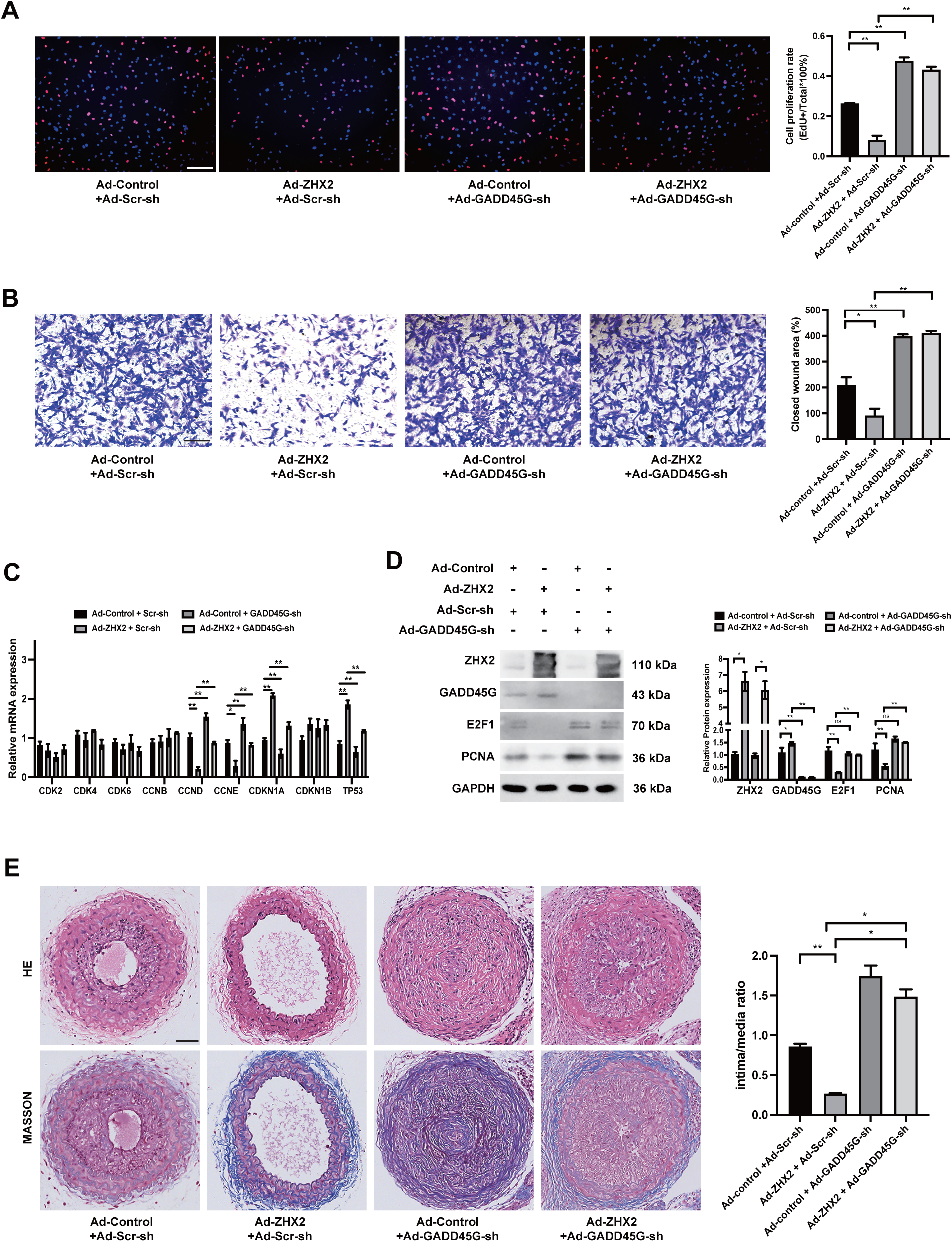
**Knockdown of GADD45G counteracts the inhibitory effect of ZHX2 overexpression on Vascular Smooth Muscle Cells (VSMCs)** (A) Examination of the proliferative capacity of *in vitro* cultured smooth muscle cells under conditions of ZHX2 overexpression and/or GADD45G knockdown, as determined by 5- Ethynyl-2’-deoxyuridine (EdU) assay. (n=3). Scale bar = 100 μm. (B) Assessment of the migratory ability of *in vitro* cultured smooth muscle cells under conditions of ZHX2 overexpression and/or GADD45G knockdown, as evaluated by Transwell migration assay. (n=3). Scale bar = 100 μm. (C, D) Determination of the mRNA or protein expression levels of ZHX2, GADD45G, and proliferation-related genes in *in vitro* cultured smooth muscle cells under conditions of ZHX2 overexpression and/or GADD45G knockdown, as measured by Western blot and Real-Time Quantitative Polymerase Chain Reaction (RT-qPCR). (n=3) (E) Evaluation of ligation-induced neointima formation under conditions of ZHX2 overexpression and/or GADD45G knockdown. (n=3). Scale bar = 50 μm. *\*p* < 0.05, *\*\*p* < 0.01, statistical significance determined by One-way Analysis of Variance (ANOVA) with Bonferroni’s Post Hoc Test or Welch’s ANOVA with Dunnett’s T3 Post-Hoc Tests.

## Discussion

Intimal hyperplasia, characterized by the accumulation of VSMC in the arterial wall, is a common feature of pathological arterial remodeling. Combatting it is crucial in life-threatening cardiovascular diseases, including coronary artery disease, stenosis and atherosclerosis. Here, we document the association of ZHX2 with intimal hyperplasia, as well as VSMC proliferation and migration, providing new insights into the discovery of therapeutic targets for vascular diseases. This study found that the expression of ZHX2 was reduced in injured arteries and in VSMCs stimulated by PDGF-bb. This suggests that ZHX2 might have an inhibitory effect on the pathology of VSMCs and adverse arterial remodeling caused by injury response. The study also shows that ZHX2 regulates GADD45G gene expression in VSMCs, which results in the inhibition of vascular smooth muscle proliferation, migration, and intimal hyperplasia. These findings provide new insights into the development of vascular diseases and may lead to the creation of innovative treatments for the prevention of such diseases.

GWAS have yielded many results in the study of cardiovascular diseases, such as common genetic variants that increase the risk of coronary artery disease (CAD) (31). There are still many genes involved in GWAS variation that have not been fully studied, and their in-depth exploration may reveal new biological mechanisms. Several genome-wide association studies have shown a strong association between changes in the ZHX2 locus and right carotid artery IMT (28-30); This suggests that ZHX2 may play an important role in pathological vascular remodeling and neointimal formation in diseases such as stenosis and atherosclerosis. We found that the expression of ZHX2 was significantly downregulated in the injury model. We looked for expression profiling data for atherosclerosis and got similar results. After arterial injury, VSMC was the main cell type in neointima. We further performed immunofluorescence analysis and found that the expression level of ZHX2 was significantly decreased in αSMA-labeled VSMCs in the injury group compared with the control group. We then constructed a PDGF-bb-stimulated *in vitro* model and found that the expression of ZHX2 decreased after PDGF-bb stimulation. These results indicated that ZHX2 gene expression in VSMCs was significantly regulated by injury stimuli, which led us to focus its function on VSMCs. However, ZHX2 may also be involved in vascular remodeling in other cell types, including but not limited to endothelial cells, fibroblasts, and/or immune cells. In order to rule out the interference of other cell types, we used vascular smooth muscle-specific ZHX2 knockout mice for subsequent ligation-induced carotid artery injury experiments. However, this does not suggest that we consider ZHX2 in other cell types to be independent of vascular remodeling and neointimal neogenesis. Conversely, ZHX2 deficiency in mouse bone marrow has been shown to ameliorate atherosclerosis by inhibiting inflammation and macrophage apoptosis (32). Just like the Arnus phenomenon named after the Roman double-faced god Arnus, the duality of ZHX2 in bone marrow cells and smooth muscle cells also reminds us that in the study of genes and vascular remoling-related diseases, the role of genes cannot be simple. The division into good or bad requires caution when intervening in the individual process. Therefore, it is necessary to specify the cell type of VSMC and understand the specific role of ZHX2 in detail because this will become a key link in the design of targeted therapeutic measures in the future. At the same time, the role and mechanism of ZHX2 in different tissues or cell types need to be further studied in the future to provide a more accurate and comprehensive scientific basis for targeted therapy.

More and more evidence shows that ZHX2 is an important regulator of various biological processes, including regulating the development of various cancers such as liver cancer, gastric cancer, and thyroid cancer, and participating in lipid metabolism, cell differentiation, and development, NK cell maturation, and macrophage cell polarization, etc (33). Many literatures have reported that ZHX2 has an inhibitory effect on the proliferation and migration of different types of cells (17, 21, 34-39). Some literature reported that ZHX2 may be related to apoptosis (35, 37). There are also some literature reports that ZHX2 overexpression can promote proliferation and migration and inhibit apoptosis (40-42). Studies have shown that ZHX2 improves atherosclerosis by regulating macrophage apoptosis. Looking ahead, we provide new evidence for the role of ZHX2 in vascular disease, suggesting that it also plays an important role in VSMC pathology. We found that overexpression of ZHX2 had no significant effect on VSMC apoptosis (Figure S4). Meanwhile, our data from vascular smooth muscle-specific knockout of ZHX2 showed that knockout of ZHX2 significantly promoted the proliferation and migration of VSMCs induced by PDGF-bb stimulation, as well as ligation-induced carotid intima neogenesis. Correspondingly, by overexpressing ZHX2, we found that it significantly reduced VSMC proliferation and inhibited neointima formation 14 days after injury. In order to fully understand the effect of ZHX2 on VSMC, we performed RNA-seq and found that overexpression of ZHX2 significantly affected the expression of VSMC cell cycle-related pathway genes. This validates the results of our experiments. These results suggest that ZHX2 is a potential therapeutic target for neointima formation, restenosis, and atherosclerosis at the core of VSMC proliferation and migration.

A significant finding of this study is the discovery that GADD45G is a downstream target of ZHX2 function in VSMC pathology. ZHX2 is a ubiquitous transcription factor belonging to the Zinc fingers and homeoboxes (ZHX) family, whose ZNF domain has DNA-binding activity. It was initially identified as a negative regulator of postnatal AFP in mice (20). The mechanism by which ZHX2 controls the transcription of target genes is not yet fully understood. To identify genes that may be regulated by ZHX2 on transcription level, we combined the results of a ChIP-seq with RNA-seq results. We obtained a total of 22 promoter regions that may be regulated by ZHX2. We annotated these genes and discovered that GADD45G is closely related to cell cycle progression. Our findings suggest that GADD45G is a transcriptional target of ZHX2 in the VSMC background. The overexpression of ZHX2 significantly up-regulated the RNA and protein expression levels of GADD45G. ChIP-qPCR experiments also verified the combination of ZHX2 and the GADD45G promoter region.

The GADD45 family proteins, including GADD45A, GADD45B, and GADD45G, are small proteins (about 18 kD) with high homology found in the nucleus and cytoplasm. They are known as "Stress Sensor Genes" as they respond to various stress signals, including DNA damage, ultraviolet radiation, oxidative stress, and chemical exposure. GADD45 family proteins have significant functional similarities and regulate several biological processes. They interact with various crucial proteins, bind RNA to affect their stability, and demethylate specific genes. These processes include cell cycle arrest, DNA repair, apoptosis, and cell survival and senescence (43, 44). GADD45G affects the cell cycle in several ways. It can inhibit the kinase activity of Cdk1/cyclinB1 by directly interacting with the CDK1/cyclinB1/PCNA complex (45). It can also induce CDKN1A’s expression (46) and downregulate the expression of E2F1 by passing the p38 MAPK pathway (47). Our research found that ZHX2’s overexpression significantly increased CDKN1A’s expression, decreased E2F1’s expression, and inhibited the proliferation and migration of VSMCs in PDGF-bb-stimulated VSMCs. Knocking down GADD45G on this basis restored the effect of ZHX2 overexpression on them. This indicates that GADD45G is the key pathway for ZHX2 to exert its inhibitory effect on VSMCs’ proliferation and migration.

Taken together, we have established the crucial role of ZHX2 in pathological arterial remodeling. Furthermore, we have identified GADD45G as the downstream target gene that is responsible for the effects of ZHX2 mediating these changes. Our findings significantly improve our understanding of neointimal hyperplasia and pathological arterial remodeling and offer potential therapeutic targets in the ZHX2/GADD45G signaling pathway for cardiovascular diseases.

## Author contributions

Z. Zheng, Y. Li and K. Huang conceived and designed the research; S. Fan, L. Yang and X. Wu performed the experiments; Y. Wu, P. Wang, and B. Qiao contributed to analysis the data; Z. Zheng, Y. Li and K. Huang wrote and revised the manuscript. All authors read and approved the final manuscript.

## Availability of data and materials

We obtained the gene expression profiles for carotid atheroma from the Gene Expression Omnibus database (GEO; http://www.ncbi.nlm.nih.gov/geo/). The accession numbers were GSE43292 and GSE70410. RNA-seq and ChIP-seq analyses were performed in this study, and these data were deposited in the NCBI Gene Expression Comprehensive database under accession number GSE244565.

## Supplementary materials

**Fig. S1.**
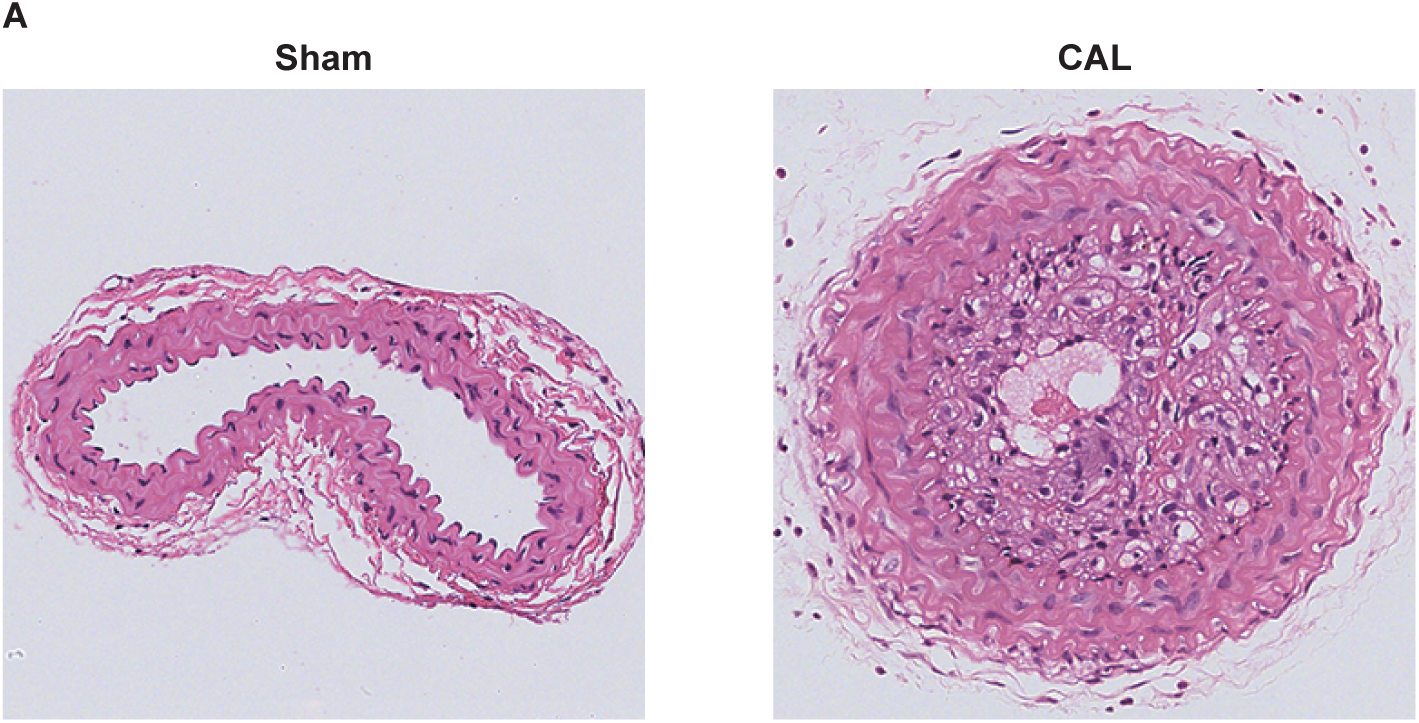
Histological Analysis of Neointimal Hyperplasia Post Carotid Artery Ligation. (Left) Hematoxylin and eosin (HE) staining of the carotid artery from the sham-operated group, showing a normal arterial structure without evident neointimal formation. (Right) HE staining of the carotid artery from the ligation group, illustrating pronounced neointimal hyperplasia post-ligation, indicative of successful model establishment. Scale bar = 50 μm.

**Fig. S2.**
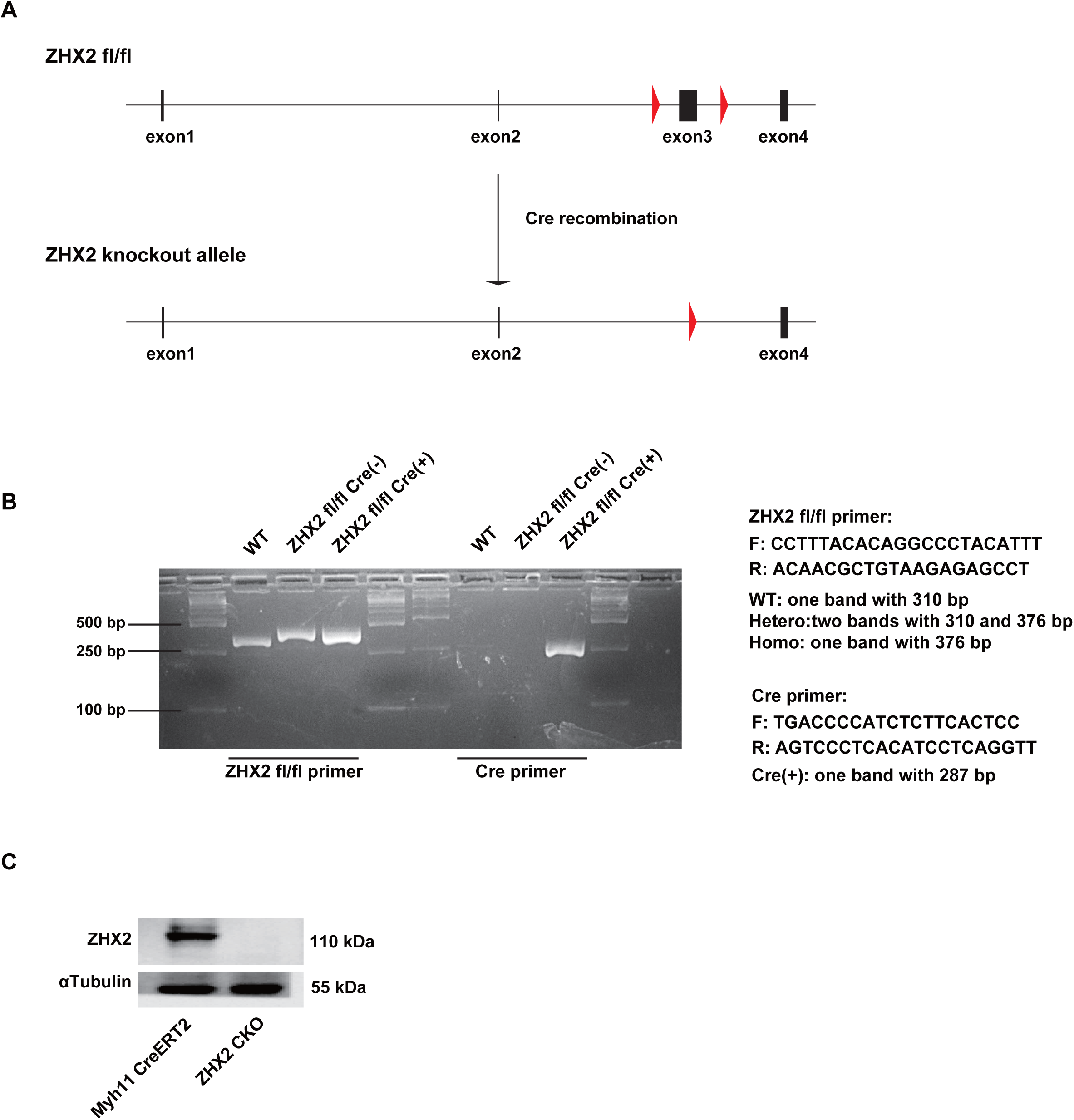
**Generation and Validation of Smooth Muscle-Specific (Myh11-CreERT2) ZHX2 Knockout Mice** (A) Schematic representation of the flox/flox construct used for creating conditional ZHX2 knockout in smooth muscle cells via Myh11-Cre-mediated recombination. (B) Agarose gel electrophoresis results for genotyping, showcasing bands corresponding to Wild-type (WT), ZHX2 fl/fl Cre(-) (non-recombined), and ZHX2 fl/fl Cre(+) (recombined) mice. (C) Western blot analysis of primary VSMCs cultured from aorta of Myh11-CreERT2 and ZHX2 CKO mice, using an antibody against ZHX2. A-Tubulin serves as the loading control.

**Fig. S3.**
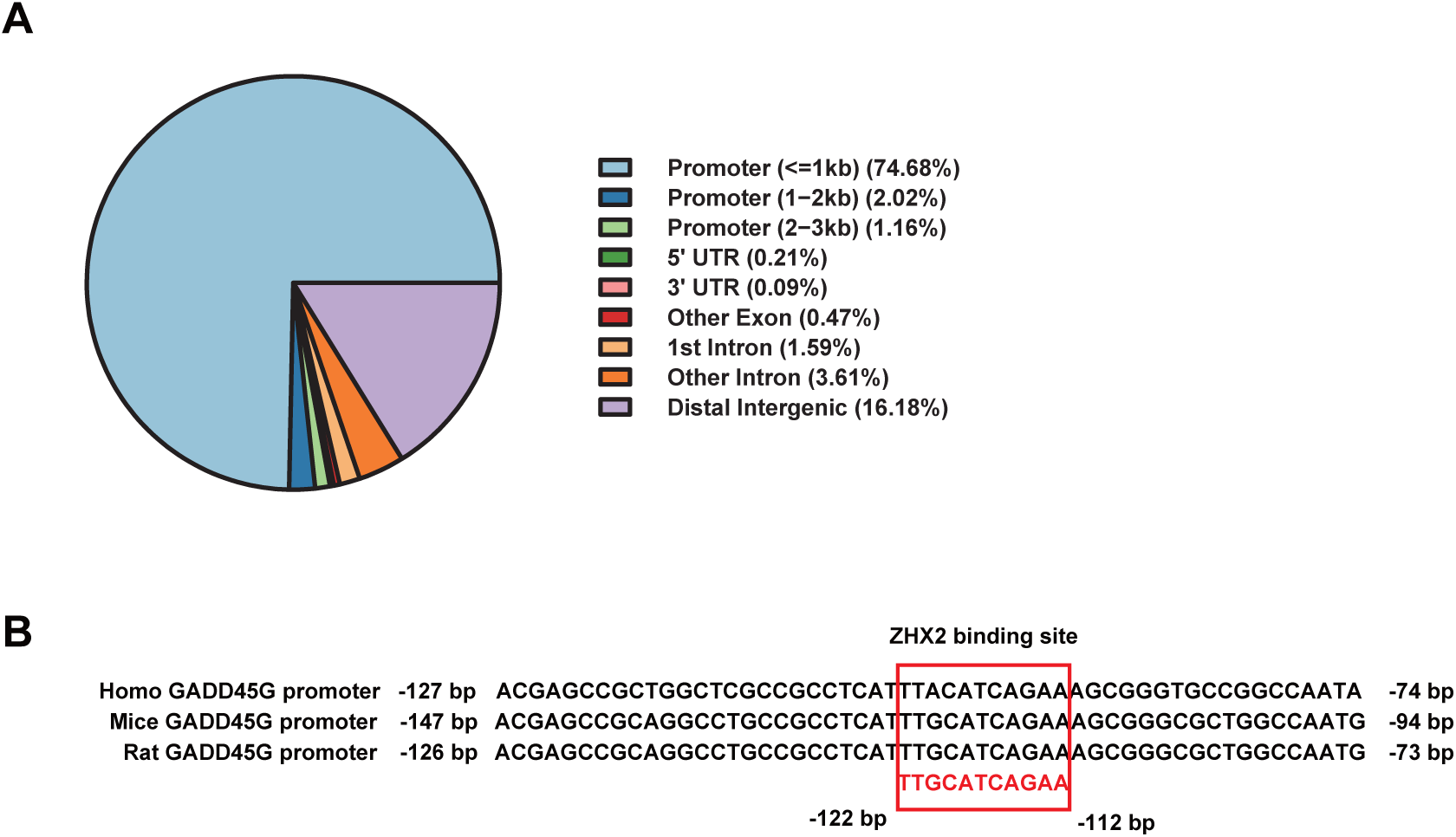
**ChIP-seq Analysis of ZHX2 Binding Patterns and Its Association with GADD45G** (A) Distribution of ZHX2 binding peaks across various genomic regions including promoters, UTRs, introns, and others. (B) Comparative analysis of the ZHX2 binding site sequences surrounding the GADD45G promoter region across human, mouse, and rat genomes, emphasizing the evolutionary conservation of ZHX2 interaction motifs.

**Fig. S4.**
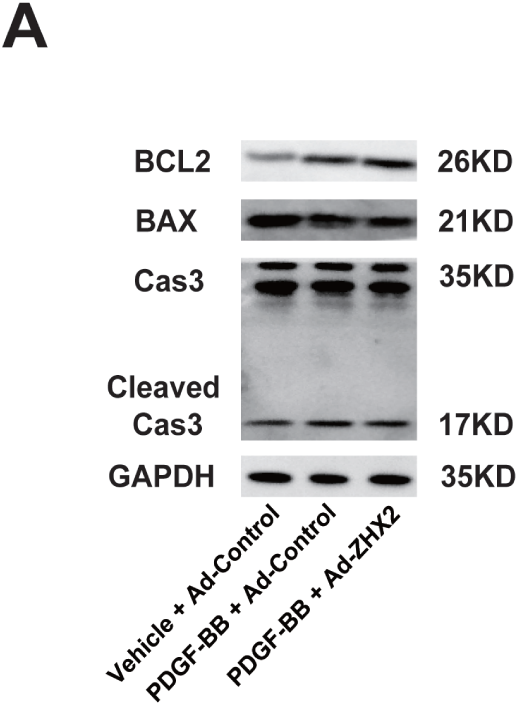
**The Impact of ZHX2 Overexpression on the Apoptosis of VSMCs** Western blot analysis illustrates the expression levels of BCL2, BAX, caspase-3 cleaver (cleaved cas3), and caspase-3 (cas3) under different conditions as shown in the figure.

**Fig. S5.**
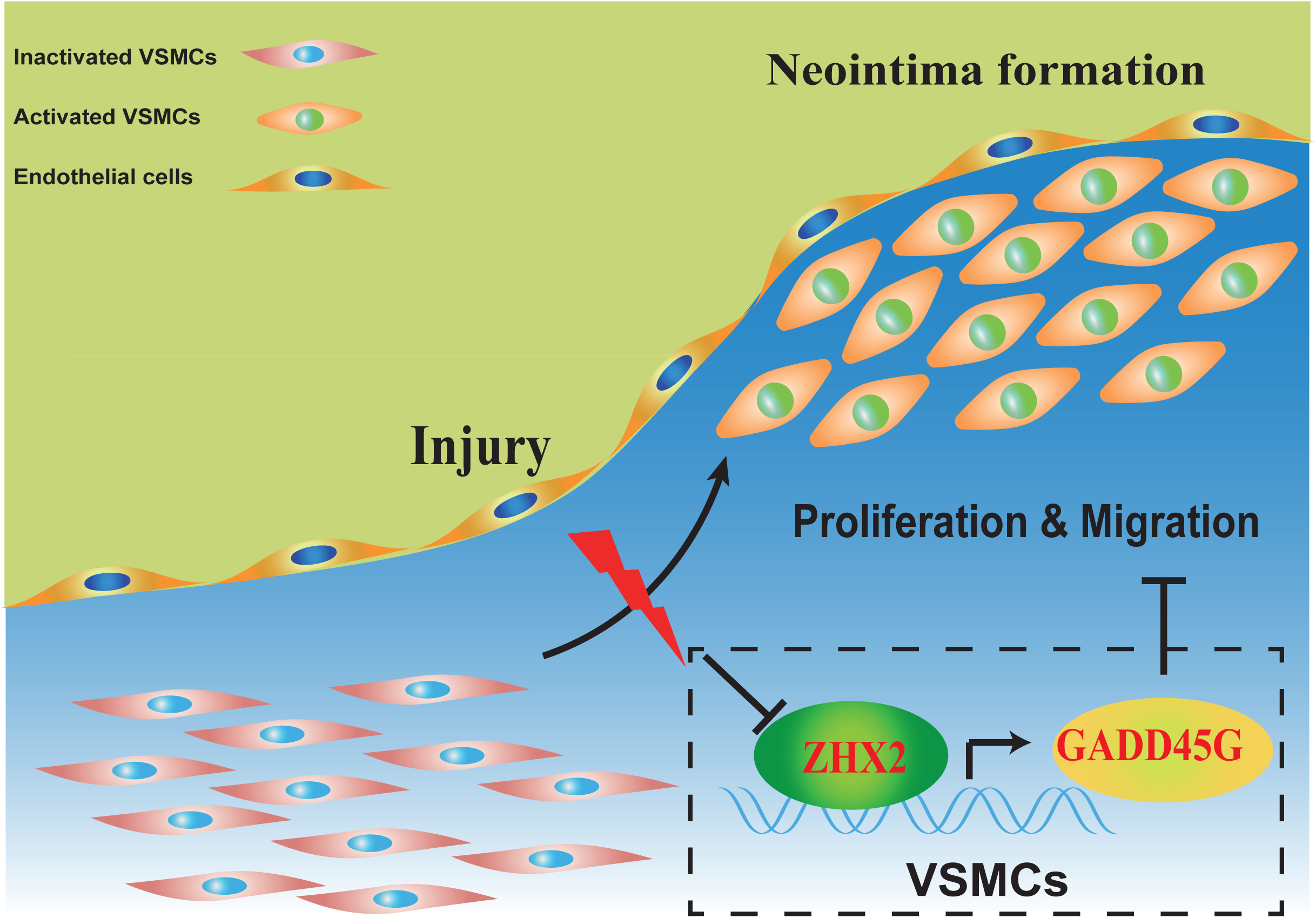
**Schematic diagram of the molecular mechanisms underlying ZHX2-regulated vascular remodeling**

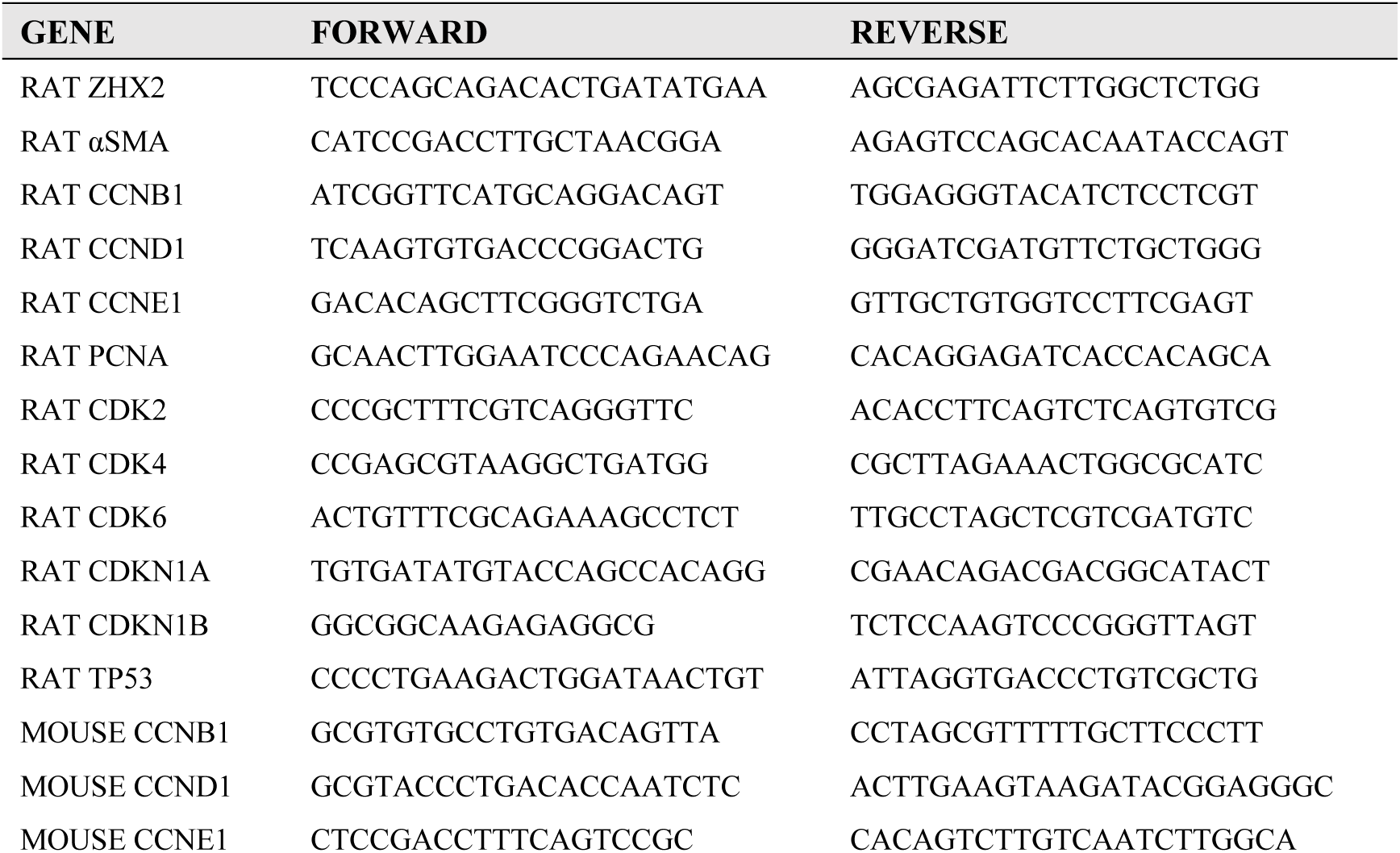
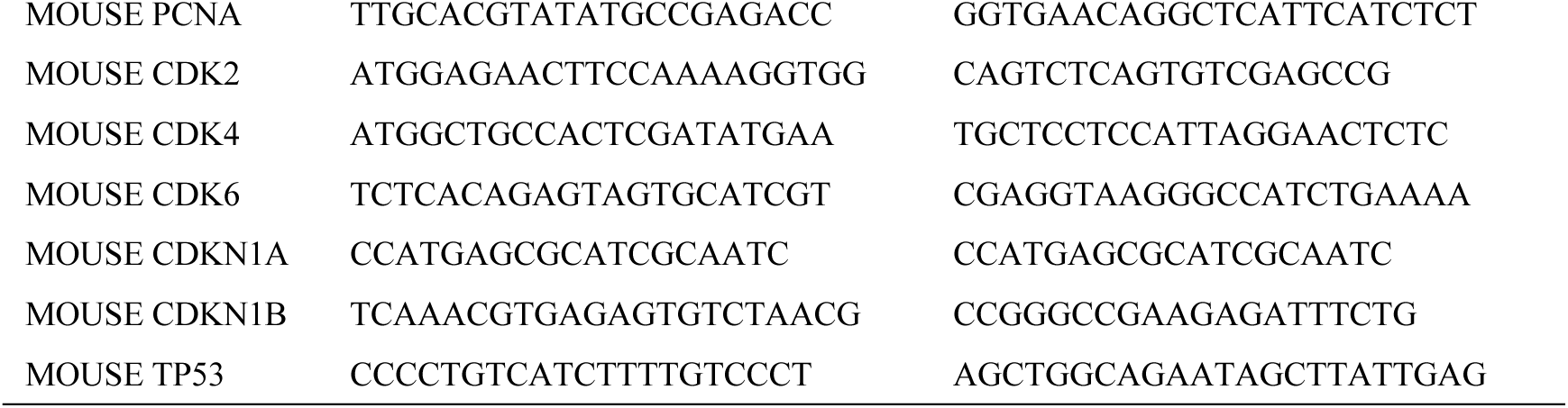
Supplementary Table 1.

